# Laser Capture Microdissection optimization for high-quality RNA in mouse brain tissue

**DOI:** 10.1101/2021.08.30.458265

**Authors:** Margareth Nogueira, Daiane CF Golbert, Richardson Leão

## Abstract

Laser Capture Microdissection (LCM) is a method that allows to select and dissecting specific structures, cell populations, or even single cells from different types of tissue to extract DNA, RNA, or proteins. It is easy to perform and precise, avoiding unwanted signals from irrelevant cells, because gene expression may be affected by a bulk of heterogeneous material in a sample. However, despite its efficiency, several steps can affect the sample RNA integrity. In comparison to DNA, RNA is a much more unstable molecule and represents a challenge in the LCM method. Here we describe an optimized protocol to provide good concentration and high-quality RNA in specific structures, such as Dentate Gyrus and CA1 in the hippocampus, basolateral amygdala and anterior cingulate cortex of mouse brain tissue.

## INTRODUCTION

In this work, we describe an optimized Laser Capture Microdissection protocol for acquiring high-quality RNA from specific structures in the mouse brain tissue. LCM method permits precise isolation of single cells or groups of cells from heterogeneous tissues to extract DNA, RNA, or proteins for further analysis through RT-qPCR, sequencing, or proteomics studies, among others. It is a powerful tool that leads to an experimental refinement, allowing more accurate results due to the collected material specificity (Sonntag and Woo, 2018; Erickson et al., 2009). This ability to acquire defined regions or cell types eliminates the transcriptional background noise of unwanted tissue leading to a localized analysis of gene expression. In a bulk, cellular heterogeneity impairs a precise understanding of gene expression, where unwanted cells may conceal the signals of the relevant cells, affecting comprehension of biological functions in physiological or pathological conditions. Thus, overcoming this limitation is a major advance in the field provided by the LCM technique (Nichterwitz et al., 2018). However, DNA analysis is more applied than RNA in LCM, because RNA is considerably a more unstable molecule, susceptible to degradation by a wide variety of RNases, both endogenous and exogenous (Mahalingam, 2018). Ribonucleases (RNase) are enzymes that catalyze the hydrolysis of RNA molecules, breaking them down into small parts. This is a common reason for failing experiments, therefore the isolation of preserved RNA is crucial for the success of the research. Tissue manipulation, LCM process, and RNA extraction have a lot of critical steps that can affect RNA integrity. RNase is present in large quantities in human skin and is difficult to inactivate. Everything that contacts human skin will be contaminated consequently and can destroy the RNA quickly, even in low amounts (Nichols et al., 2008; Morrison et al., 2012; Garrido-Gil et al., 2017; Meda et al., 2019). Due to the specificity of LCM, a small quantity of samples is acquired from each sample and isolating high-quality RNA often still represents a challenge. Therefore, to preserve RNA integrity some cautions are required (Schaeck et al., 2016).

Endogenous control of RNase is mainly done by low temperatures. Exogenous control is more complex, and depends on a wider range of actions. We have optimized critical steps and adapted the workflow of the basic LCM protocol to minimize handling, transport, and temperature variation on the samples to obtain high-quality RNA for gene expression analysis in different structures, as the hippocampus, amygdala, and anterior cingulate cortex from mouse brain sections. It is important to highlight that the LCM protocol presented here allows the analysis of gene expression in several mammalian brain structures and not only in adult mice but also in embryonic stages. In order to validate this new protocol to reach a good quality of the LCM-derived RNA, we performed three independent experiments. We analyzed the BDNF gene expression in three different regions of the mouse brain with RT-qPCR ranging from one hour, five hours, and 5 days after two different treatments. The result of our quality control presented a total of 94% of high-quality RNA in 70 samples, obtained under a bioanalyzer device. 18 samples obtained RIN = 10, totalizing 26%. Therefore, under specific conditions and high performance, we conclude that our optimized protocol can be useful for experiments using microdissected mouse brain tissue, resulting in RNA with a high degree of purity.

## STRATEGIC PLANNING

Before starting the acquisition of samples, there are many solutions, material decontamination, brain dissections, and LCM settings to be prepared. All of this work can be done previously or a day in advance to avoid work overload, since cryosectioning, staining, and microdissection are time-consuming processes that are required to be performed carefully to prevent any contamination and RNA degradation. Our workflow starts with cleansing RNase free, then cryosectioning followed by staining/fixation, and finally the LCM collecting sample. As soon as the first slide of tissue is ready to use, it is taken to the LMC system for microdissection and only after that, the second slide will be cut. During this period the brain stays in the cryostat, protected from RNase at −20 ° C. In other words, we only prepare the subsequent slide after collecting the previous one. This new approach avoids excessive manipulation and exposure to RNase preventing consequent RNA degradation. We work continuously for at least 8 slides, where each slide contains 6 to 8 slices. Each slide sample takes between 50 minutes and one hour to be acquired (10-15 minutes for cryosectioning/staining/dehydration and 40-45 min on LCM system). After cutting and catapulting, we transfer the isolated tissue to a collection tube inserted in an icebox. After 8 – 10 repetitions, that is, 8-10 slides with 6 – 8 slices, we spin down the amount collected and store at −80°C freezer for further RNA extraction (Fig 1).

**Figure 1.**
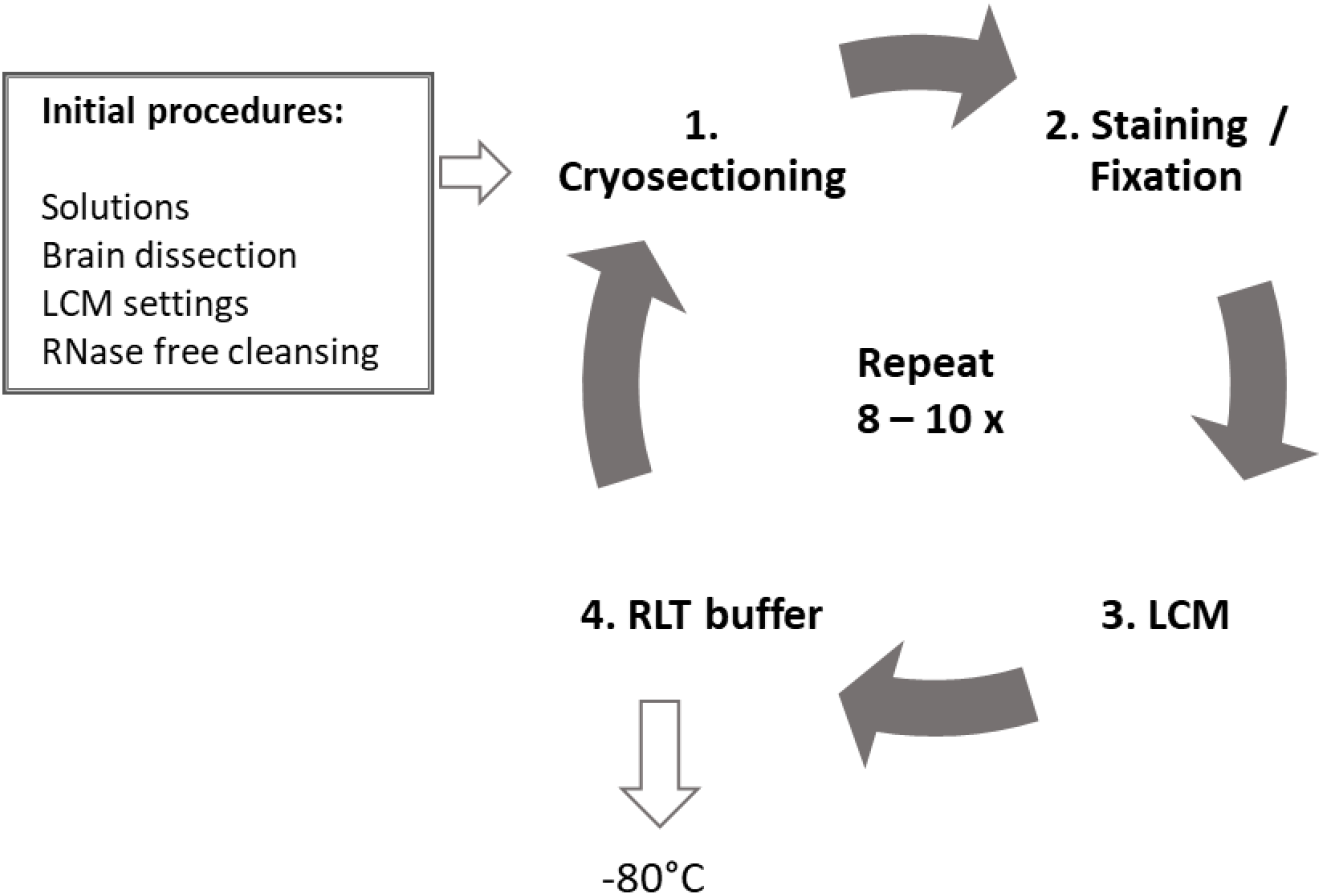
Schematic workflow for the experiment of isolating subpopulation regions from the mouse brain using the LCM technique. The initial procedures are listed as previous work in the box, and the following steps include 1. Cryosectioning, 2. Staining/Fixation, 3. LCM, 4. Transfer collected tissue to collection tube with RLT buffer, repeated from 8 to 10 times, to each sample. The collected samples are stored in the −80° C until posterior RNA extraction.

## LASER CAPTURE MICRODISSECTION OF MOUSE BRAIN TISSUE

The following protocol for LCM employs the PALM MicroBeam laser microdissection and catapulting system (Fig 2A). Although the procedure collected dentate gyrus, CA1, amygdala, and anterior cingulate cortex tissue, this method can be used for a variety of regions in brain tissue, without modifications. It is not applicable for Formalin-Fixed Paraffin-Embedded (FFPE) sections.

**Figure 2.**
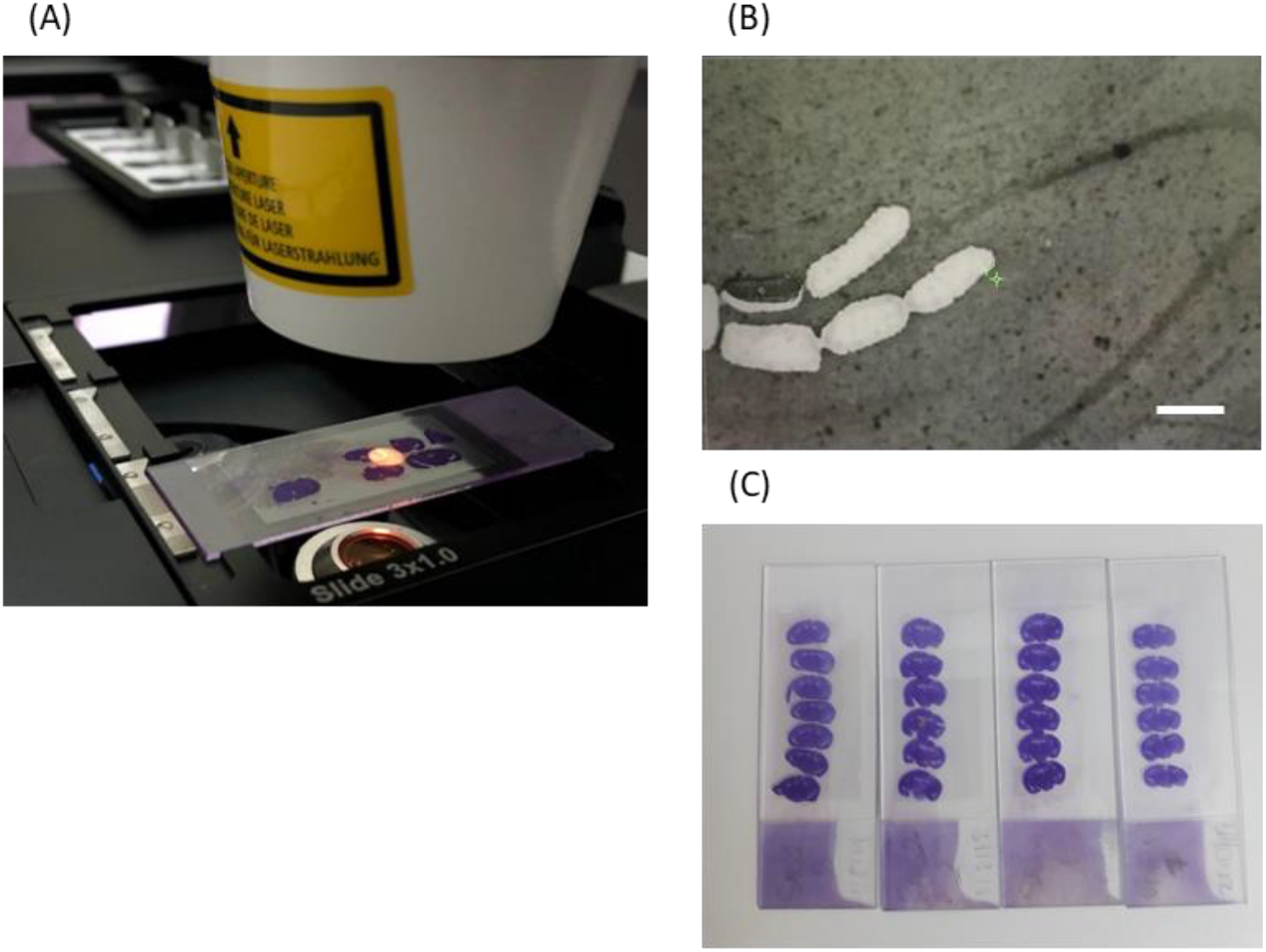
LCM features and staining. **(A-B)** PALM microbeam setup in action and screen image. Scale bar: 200 μm. **(C)** Examples of violet cresyl staining slide with six or seven brain slices of mouse brain tissue.

### Materials List

#### Materials

1. PEN Membrane Slides (P.A.L.M.microbeam)
2. PALM adhesive cap tubes (Carl Zeiss)
3. RNeasy Micro kit (QIAGEN)
4. Cryostat blade
5. Falcon tube RNase free 50mL
6. Pipettes
7. Filtered tips RNase free
8. RNase free tips
9. Tubes RNase free (Eppendorf)
10. Amber bottle
11. Gloves and mask
12. Needles
13. Parafilm
14. Aluminum foil
15. Pen
16. Pencil
17. Storage box
18. RNase free wipes
19. Paper towels
20. Paintbrush (fine point)
21. Scissor
22. Tweezer
23. Single edge Razor Blade

#### Solutions

1. Violet Cresyl Acetate Solution (1% w/v)
2. Water DEPC 0.01%
3. Water DEPC 0. 1%
4. PBS DEPC
5. 50% RNase free ethanol
6. 70% RNase free ethanol
7. 100% RNase free ethanol
8. RLT buffer

#### Equipment

1. PALM MicroBeam System (Carl Zeiss)
2. Computer
3. PALM RoboSoftware (Zeiss)
4. Cryostat
5. Fume hood
6. Microcentrifuge MiniSpin (Eppendorf)
7. Autoclave and Lab oven
8. Thermocycler
9. ABI ViiA 7 Real-Time PCR System (Applied Biosystems, NY, USA)
10. Bioanalyzer (Agilent)
11. Bioanalyzer RNA 6000 Nano assay
12. ND8000 spectrophotometer (Thermo Scientific NanoDrop Products)
13. Freezer −80° C
14. Fridge
15. Timer

### Initial procedures

a. Preparing solutions: *see recipes below*
b. Cleaning PEN slides
  1. Wear gloves, lab coats and a cap.
  2. Wipe the PEN slides with RNase-Zap, wash with DEPC water and then put to UV (ultra-violet) dry under the fume hood for 30 minutes, or overnight at 200° C in the bath or at 180° C in the oven for 4 hours.
  *Put the slides in the UV for 30 minutes each time they will be used, even if they are previously cleaned*. *The cleansing process is crucial for the maintenance of RNA integrity. Wear gloves all the time and change them frequently. Cleaning the gloves with RNase-Zap wipes is a good practice either. Before starting any procedure involving tissue samples, wipe all the instruments and equipment with 70% ethanol RNase free (with DEPC water 0,01%) and RNase-Zap (Sigma). Use certified RNase-free materials whenever is possible and be sure they are well cleaned. Avoid handling the samples when is not necessary to diminish contamination and RNA degradation* (Nichterwitz et al., 2018). *Store all materials and reagents for RNA separate from others*. *RNase-Zap must be used sparingly, otherwise the excess may affect the sample purity*.
c. Dissection and storage of the brain
  3. Wear gloves, lab coats and a cap.
  4. Clean workbenches and tools with RNase free 70% ethanol. Wipe RNase-Zap on all the surfaces and materials that contact directly the sample.
  5. Deeply anesthetize the mouse.
  6. Decapitate and quickly dissect the brain. The dissecting process must be done in approximately 2 minutes to maintain the integrity of RNA.
  7. Remove the blood excess with PBS RNase free. This step is important due to the large amount of RNase found on blood.
  8. Submerge the brain in isopentane at −40 ° C for 30 seconds. Use a styrofoam box with dry ice for transportation, if necessary.
  9. Wrap the brain in a clean aluminum foil and store on −80 ° C freezer until cryosectioning.
d. LCM settings We performed the LCM calibration test using the PALM RoboSoftware, from Zeiss. For cut energy, the optimal value we found was 50 and for laser pressure catapult (LPC) energy the best value was 80. During adjustment always keep the LPC energy higher than cut energy. We used magnification (5x) to draw the region of interest (Fig 2B). Before cutting and catapulting always set up focus and energy in a piece of tissue outside your ROI. As the cell density and the extracellular matrix can vary according to the age of the animal and the brain region, it is important to highlight that these values may vary among tissues or staining procedures, so they can be used just like a first reference, and sometimes they have to be adjusted.

### Cryosectioning

10. Wear gloves, coats and a cap.
11. Clean cryostat and tools with RNase free 70% ethanol. Wipe RNase-Zap on all the surfaces and materials that contact directly the sample, like brush tips and cutting blade.
12. Stabilize the brain on −20° C inside the cryostat, for 30 min.
13. Organize materials for staining to keep them cold (see in staining and fixation section).
14. Number and label the slides with a pencil, pen dye may not work at cold and in stained slides.
15. Lay down some slides inside the cryostat to maintain them cold. We put in groups of three.
16. Cut 12μm slices and at the moment of transferring the sections to the slide, heat a lower part of the slide with the finger underneath the slices to favor adhesion to the cut.
17. Wait for the slide to dry inside the cryostat for 2-3 min.

*Cut between 6 and 8 slices in a slide, the optimal time is between 10-15min to prepare each slide. 30 – 45 min is the optimal time in the LCM system. Slides must not take more than one hour at room temperature, otherwise, RNA quality may be compromised as endogenous RNases may still be active*.

*We strongly suggest turning on all the LCM equipment before cryosectioning to be sure that everything is working fine*.

*Avoid embedding the whole brain in OCT because it can cause some damage to RNA integrity. This freezing medium interferes with laser efficiency. In this protocol, we use a few amounts of OCT just to fix the brain in the cryostat where it will be at −20*^*o*^ *C until the end of the work*.

*We performed thickness tests, with 10µm, 12µm and 16µm. Adult mouse brain hippocampus, and amygdala tissue presented better results with 12µm thickness. These values can vary among structures due to different patterns of extracellular matrix and cell density*.

### Staining and fixation

18. After drying in the cryostat, soak the slide in 70% ethanol RNase free (falcon tube inserted in an icebox) for 2 min.
19. Soak the slide in violet cresyl for 30 seconds (falcon tube inserted in icebox).
20. Remove the excess of violet cresyl with paper towels.
21. Dip the slide quickly in 70% ethanol RNase free (falcon tube inserted in icebox).
22. Dip the slide quickly in 100% ethanol RNase free (falcon tube inserted in icebox).
23. Allow it drying at room temperature for 1-2 minutes.
24. Use immediately on LCM for no more than 45 min. This precaution is important for the maintenance of RNA integrity.

*We followed the manufacturer orientations for staining (PALM Protocols – RNA handling, from Carl Zeiss)*

*Fill a big styrofoam box with ice. Organize a 50ml falcon tube with 70% ethanol Rnase free, then a 50ml falcon tube with violet cresyl, a 50ml falcon tube with 70% ethanol Rnase free to be used after violet cresyl and a 50ml falcon tube with 100% ethanol Rnase free. Let some paper towels beside the box to clean the violet cresyl excess. Control the time with a timer*.

*In our experience, fixation is crucial to ensure a high-quality RNA and any failure can degrade the RNA. Be sure to have clean gloves and do not touch anything unnecessarily*.

*Proper dehydration of tissue slices minimizes upward adhesive force between the slide and the tissue (Datta et al., 2015)*.

*Violet cresyl shows good visualization and absorbs the laser energy, preventing damage to the cellular components (Chabrat et al., 2015; Mahalingam, 2018). In our experience, it was a good choice (Fig 2C)*.

*High temperature degrades RNA. When the tissue is out of cryostat, at room temperature, all the solutions must be inserted on ice, to keep them cold. RNA is quickly degraded, so it requires stringent RNase-free sterile conditions during handling and preparation with, in some cases, additional use of commercially available RNase inhibitors*.

### Slide storage

It is important to highlight that there is no storage before LCM in our protocol, we do not use previously frozen slides. The brain stays in the cryostat at −20° C. We prepare the slide and collect the sample in sequence, one by one, as explained earlier. Time and temperature are crucial to maintaining the integrity of RNA and we consider this step fundamental to the process.

### Laser Capture Microdissection (LCM)

We used a PALM MicroBeam system (PALM MicroBeam). The system utilizes an inverted microscope with a focused laser beam to cut out and catapult precisely selected areas without any contact. The microdissection process was visualized with an AxioCam camera coupled to a computer and controlled by a PALM RoboSoftware, that controls a motorized stage (Garrido-Gil et al., 2017).

25. Turn on the LCM system.
26. Proceed calibration test, following manufacturer instructions. Define the best settings of cutting and catapulting energies;
27. Clean workbenches and tools with RNase-free 70% ethanol.
28. 28.Place microtubes fixing the cap facing down at the LCM system.
29. Prepare an icebox and insert an RNase-free tube to receive collected samples. In a previous clean area, place a 10µl pipette, tip box, RLT buffer, and a needle.
30. After making the first slide, pin it on the LCM system and chose the working stage. Place slides into the slide holder with the membrane facing up.
31. Adjust the timer to 30 min.
32. Start drawing the region of interest (ROI). Search for cells of interest at 5x magnification. Draw cutting outlines not so close to the cells to avoid burning the tissue of interest. If there is a little distance from the cells, they can be preserved. Observe that we did not work with single cells where the procedure adopted could be different.
33. Cutting and catapulting (Fig 3A).
34. Transfer the isolated material into the sticky cap tube to the collection tube. Add 5 µl of RLT buffer into the sticky cap tube, pull the tissue with the pipette taking care to not pull too much, and transfer catapulted sections from the adhesive microtube cap to the RNase free collection tube. When enough ROI has been collected, inspect the collector and unload the tube holder (Fig 3B, 3C).
35. Insert the collection tube with sample and RLT buffer in the icebox.
36. Go to the cryostat and proceed to the subsequent slide.
37. Keep this process until the eighth slide.
38. After finishing the acquisition, spin down the collection tube in a microcentrifuge for 15 seconds.
39. Seal collection tube with Parafilm. Store the collection tube at −80 ° C freezer until RNA extraction.

**Figure 3.**
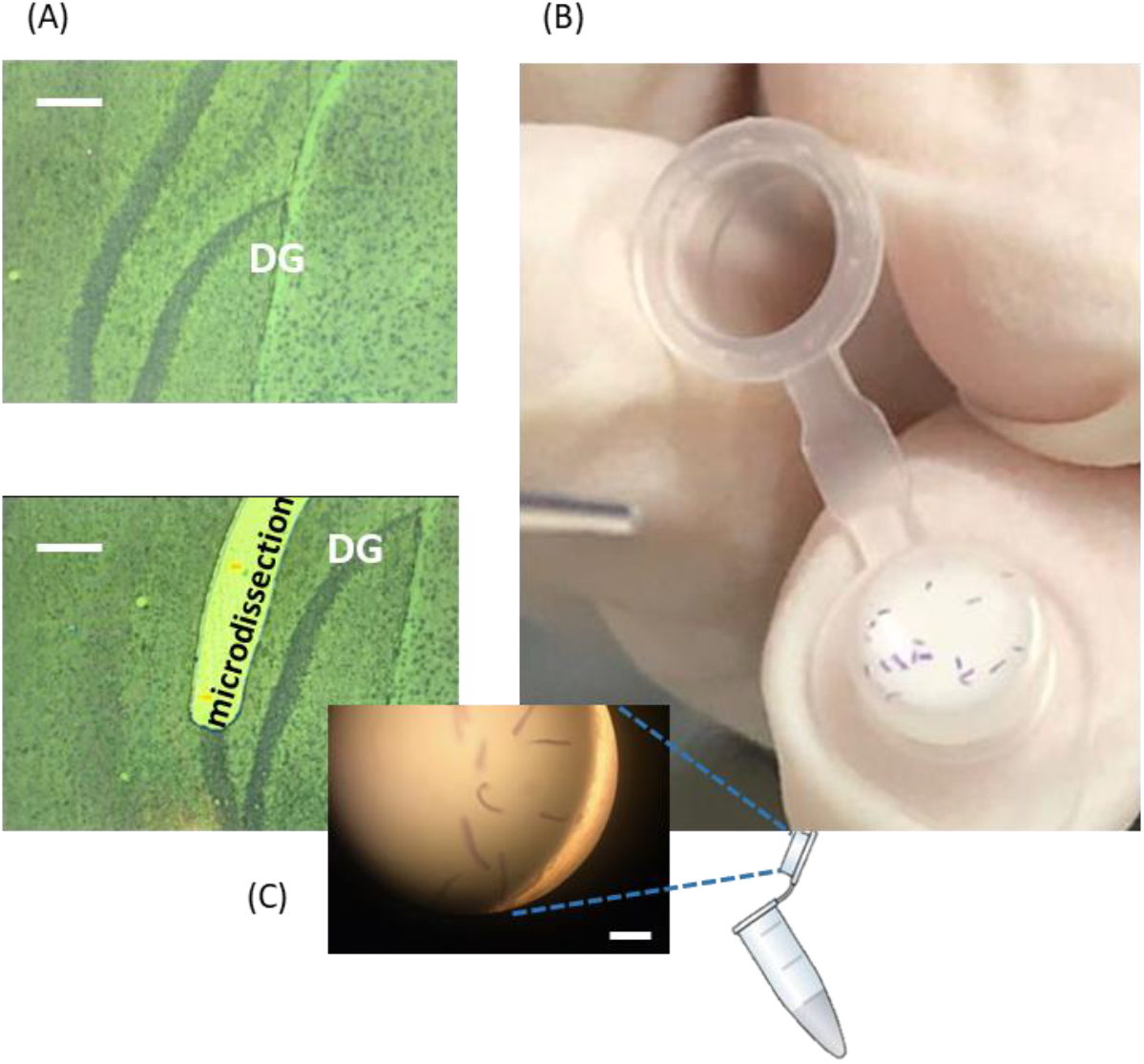
Laser Capture Microdissection process. **(A)** Dentate Gyrus before (top picture) and after (bottom picture) microdissection. Scale bar: 200 μm. **(B)** Collection of microdissected tissue from the sticky cap tube to the collection tube containg RLT buffer. **(C)** Amplified image of sticky cap tube containing Dentate Gyrus collected small pieces. Scale bar: 800 μm.

*Do the entire procedure within 30 - 45min*.

*If the material is microdissected but not catapulted, we used a previous RNase-Zap cleaned needle to remove the tissue from the slide and transfer it to the collection tube*.

*Samples were stored in an RLT buffer at −80 ° C for up to one month*.

*We made two types of collection, small pieces with a high probability of catapulting success and large extensions with a low probability of catapulting. In these cases, not catapulted laser-cut areas were removed from the slide with the slight touch of a needle to which they stuck, and then we transferred to the RLT buffer tube. In our experience, this second type of collection presented samples with higher concentrations, although it took longer. It is important to highlight that this conclusion is based on mouse hippocampus and amygdala tissue. Other types of tissues may present other patterns of RNA concentration due to differences in the extracellular matrix and cellular density, for instance. Thus, larges pieces may not configure a pattern of high-quality RNA control for all tissues*.

## REAGENTS AND SOLUTIONS

### Reagents

1. 96-100% Ethanol. Keep the container tightly closed. Keep away from heat / spark / open flame / hot surfaces.
2. RNaseZap (Ambion, #9780). Store at room temperature.
3. Tissue-Tek® O.C.T.™ (Sakura). Store at room temperature.
4. Diethyl Pyrocarbonate (DEPC) (Sigma). Keep the container tightly closed. Store at 2 - 8°C.
5. Milli-Q water.
6. Violet Cresyl Vetec V 000 178 (Aldrich). Store at 15 - 30° C.
7. β–Mercaptoethanol (Sigma). Keep container tightly closed in a dry and well-ventilated place. Containers that are opened must be carefully resealed and kept upright to prevent leakage. Store at 2 −8 °C.
8. Isopentane. Store below 50°C. Keep out of direct sunlight. May be stored under nitrogen.
9. PBS tablet (Sigma-Aldrich). Store at room temperature.
10. Ice, dry ice. Store in the freezer.
11. RNeasy Micro kit (Qiagen). Store the RNeasy MinElute spin columns and the RNase-Free DNase Set (i.e., the box containing RNase-free DNase, Buffer RDD and RNase-free water) immediately upon receipt at 2–8°C. Store the remaining components of the kit dry at room temperature (15–25°C).
12. SyBR Green/ROX PCR Mix (QIAGEN, CA, USA). SYBR Green PCR Kit should be stored immediately upon receipt at –20°C in a constant-temperature freezer and protected from light.
13. SuperScript IV Reverse Transcriptase (ThermoFisher Scientific, NY, USA). Store at the fridge.

### Solutions

a. **Water DEPC 0.01%:** Store at room temperature in an amber bottle protected from light.
  1. Add 100µl DEPC to 1L Milli-Q water, shake and leave at 37° C overnight.
  2. Autoclaving the next day. Better for washing slides and for making alcohol.
b. **Water DEPC 0.1%:** Store at room temperature in an amber bottle protected from light.
  1. Add 1mL DEPC to 1L Milli-Q water, shake and leave at 37° C overnight.
  2. Autoclaving the next day. Better for direct sample contact.
  *Do all the procedures under the fume hood. DEPC is toxic before autoclaving. Water nuclease-free is a DEPC water substitute but is more expensive. Water DEPC inhibits only RNase, while nuclease-free water inhibits RNase and nuclease either*.
c. **PBS DEPC:** Store at room temperature in an amber bottle protected from light.
  1. Dissolve a PBS RNase free pellet in 200 mL of **0.01%** DEPC water.
d. **70% RNase free ethanol:** Store in the fridge.
  1. Add 35ml of 100% ethanol and 15ml of **0**,**01%** DEPC water.
  2. Store in a 50mL falcon tube for cleaning.
e. **70% RNase free ethanol:** Store in the fridge.
  1. Add 70ml of 100% ethanol and 30ml of **0**,**1%** DEPC water.
  2. Store in two 50mL falcon tubes for the fixation process.
f. **50% RNase free ethanol:** Store in the fridge.
  1. Add 50ml of 100% ethanol and 50ml of **0.1**% DEPC water.
  2. Store for preparing the violet cresyl solution.
g. **Violet Cresyl Acetate Solution (1% w/v):** Store in the fridge in an amber bottle protected from light.
  1. Dissolve 1g of solid violet cresyl acetate in 100ml of 50% RNase free ethanol at room temperature with agitation overnight in the dark.
  *Prepare all the solutions one day in advance before to start acquiring the samples*.
h. **RLT buffer:** Store at room temperature.
  1. Add 10µl Beta-Mercaptoethanol in each 1mL RLT Buffer.

## COMMENTARY

### Background Information

The combination of a laser beam to a microscope cell microsurgery with laser was first described in the early 1960s. The laser beam of LCM has evolved for years from ultraviolet (UV) to high-energy nitrogen, infrared, and carbon dioxide lasers. In the middle of the nineties, another laser microdissection technique was developed. A carbon dioxide laser for polymerization was combined with a transparent thermoplastic membrane (ethylene vinyl acetate polymer on a glass slide). This system was successfully developed and marketed by Arcturus and after by PALM Microbeam System from Zeiss and LMD system from Leica Microsystems (Gilbrich-Wille, 2013; Chung and Shen, 2015). Although several gains have been reached, the main challenge of working with RNA analysis persists. Commonly, the results are not the most satisfactory in terms of RNA integrity, so optimization for better quality in the final product is essential.

### Critical Parameters and Troubleshooting

#### Previous work

A frequent issue at the beginning of the work is not preparing solutions and LCM settings properly, previously. Time spent in each procedure influences the final result, so it is a common problem having to stop everything to set the energy of LCM or to make a solution when time is passing on. Regarding PEN slides, it is important to observe whether the slide membrane does not have bubbles, as the adhesion of the tissue can be compromised and the process of cutting and catapulting fails. Solution: if bubbles are found, switch slides immediately.

#### Laser Capture Microdissection

A common problem in LCM is the failure of cutting or catapulting phases, resulting in the material standing over the tissue. A lot of issues can be addressed to explain not removed tissue in the LCM process. The first is a lack of adherence of the tissue to the slide membrane. A solution is performing a complete and precise in-time dehydration. Another issue may be due to a low cut or LPC energy on the LCM system. LCM’s previous settings are important, but if it is not possible, a quick calibration may be necessary to adjust the energy. However, even taking these precautions, targeted tissue may not be removed. So, a needle tip or a microcapillary can be used to remove the material with a slight touch to which they stick and then can be transferred to the collection tube cap containing 5µl of RLT buffer. This procedure requires skill and practice (Mahalingam, 2018; Fend and Raffeld, 2000).

### Understanding Results

#### RNA extraction and Quality control

The concentration and integrity of RNA are crucial for gene expression analysis. Laser Capture Microdissection generally collects a low amount of RNA. Since LCM is a complex method, several steps can affect RNA yield and quality, like cutting, staining, microdissecting, and capturing (Garrido-Gil et al., 2017; Mauney et al., 2018). We optimize these critical parameters to reach better quality control. However, the RNA extraction may affect RNA quality either. There is no point in controlling the LCM process if RNA extraction is not controlled either (Cummings et al., 2018). We extracted RNA from LCM collected tissue using RNeasy® Micro Kit (QIAGEN, #74004), which includes a genomic DNA elimination step. We followed all steps according to the manufacturer’s protocol and followed indications from The MIQE guidelines (Bustin et al., 2009). Regarding the quality control, purity, and quantity of total RNA, we applied two relevant tools: ND8000 spectrophotometer (Thermo Scientific NanoDrop Products, DE, EUA) and microfluidic analysis Agilent Technologies’ Bioanalyzer (Agilent, CA, USA) (supplementary data, table 1). The absorption spectroscopy provided by NanoDrop is an important indicator of nucleic acid purity, where absorbance reading at 260 nm (A260) is quantitative for relatively pure nucleic acid preparations in microgram quantities.

Furthermore, contaminants that can be present during RNA extraction could be accessed through absorbance. The most common are protein and aromatic moieties, which have a peak absorption at 280 nm and 230 nm respectively. Absorbance ratios 260/230 and 260/280 are used to assess the purity of RNA samples after all steps up to RNA extraction, where ratios around 2 are generally accepted as “pure” for RNA (Thermo Fisher Scientific, 2010; Gallagher, 2011). Specifically, according to Thermo Fisher Scientific (2010), NanoDrop spectrophotometers works with a lower limit of 2 ng/μL per sample, if the sample presents a low concentration (as commonly verified in LCM samples) this may result in an inconclusive result, becoming not considered 260/280 and/or 260/230 ratios, as observed in some of our samples. Bioanalyzer RNA 6000 Nano assay was used to access RNA quality through the tool RIN (RNA Integrity Number) from brain samples, considering the RNA possibility of being rapidly digested in the presence of ubiquitous RNase enzyme. This classification is based on a numbering system with 1 to most degraded RNA and 10 to the most intact one. We defined the threshold for high-quality as RIN higher than 8.5. The RIN software algorithm is robust and based on a combination of different features of RNA, as the 18 and 28S ratio, provided by microcapillary electrophoresis. It was developed to classify RNA characteristics using the Bayesian approach to train and select a prediction model based on artificial neural networks (Schroeder et al., 2006; Mueller et al., 2000).

We determined the RIN of 70 mouse brain samples. Tissues were collected from the anterior cingulate cortex, BLA amygdala, dentate gyrus, and CA1 region from the hippocampus. In the first experiment, RNA analysis shows that from a total of 19 samples, 16 presented high-quality RNA (between 8.5 and 10.0), meaning that 84% of the RNA samples were highly intact and only three presented degradation (1, 7 and 18, fig 4A). However, samples 1 and 2 are from the same animal, similar to (7 and 8) and (18 and 19). 1, 7, and 18 presented bad integrity, but 2, 8, and 19 presented high-quality RNA. As the entire procedure of slides staining and LCM were done at the same time (for example, 1 refers to the dentate gyrus of the same animal than 2, which refer to the amygdala), this suggests that the observed RNA degradation is not related to the LCM protocol, but probably to posterior RNA extraction procedure, when the samples were separate in brain regions. In the second experiment, we achieved 17 from 18 samples with high-quality RNA, representing 95% of all samples (Fig 4B). Experiment 3 presented 100% of high-quality RNA from 33 samples. The RIN range was over 9.2. The RIN of 17 from 33 samples was 10, representing 51% of all samples (Fig 4C). Figure 4D shows the percentage of RIN range among all the 70 samples, as follows: 94% from 8.5 to 10, as high-quality RNA; 1% from 7 to 8.5, good-quality RNA; 0% from 5 to 7 is a regular-quality RNA, and 4% from 0 to 5, a bad-quality RNA. This classification of RIN in quality levels was considered by this group in this specific case, with mouse brain frozen tissue.

**Figure 4.**
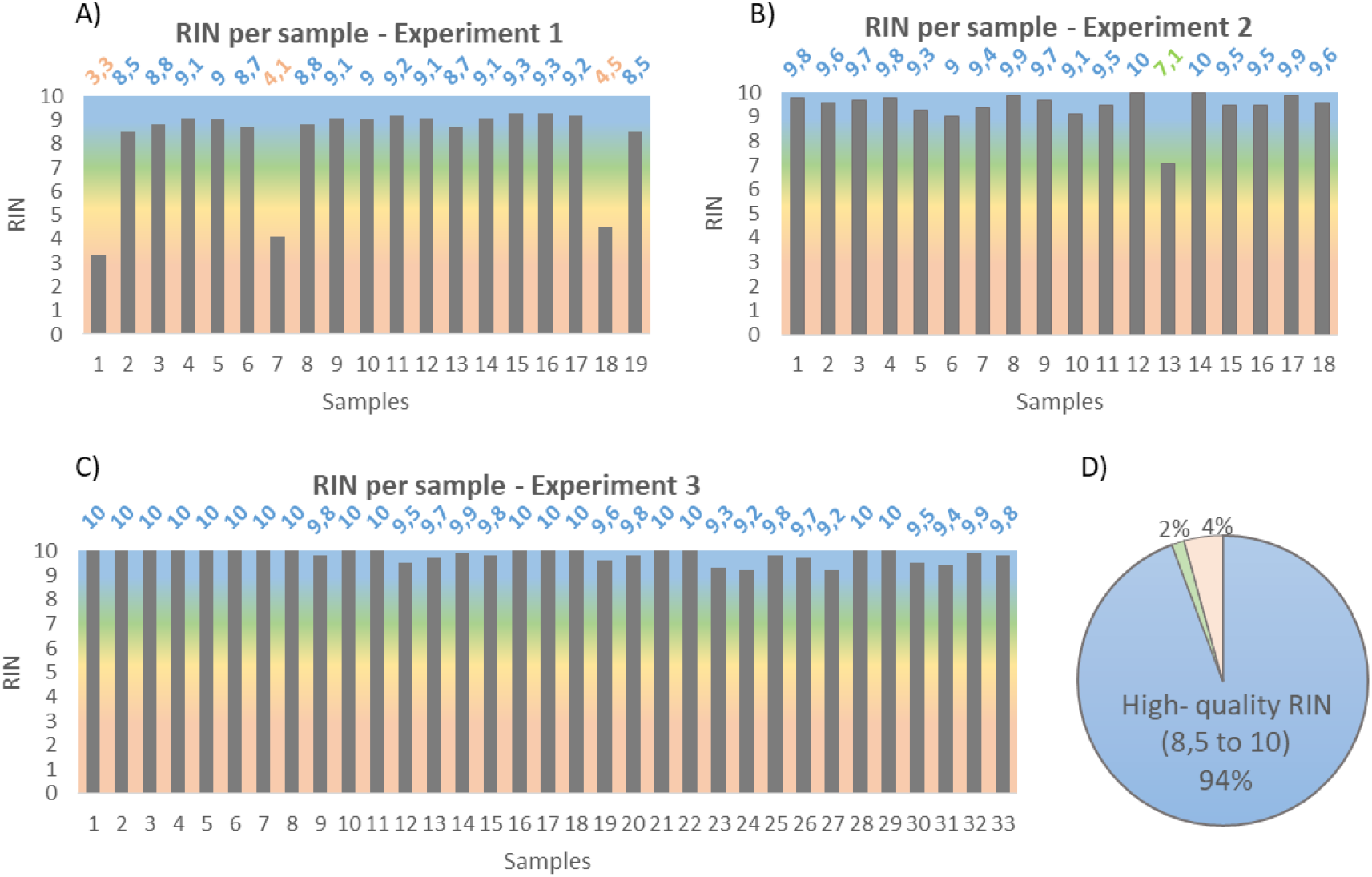
RNA Integrity Number per sample. **(A)** RIN values from experiment 1 **(B)** RIN values from experiment 2 **(C)** RIN values from experiment 3. **(D)** Pie chart showing percentage of RIN quality control from total samples. RIN below 5 is considered bad (orange), 5 - 7 regular (yellow), above 7 is good integrity (green) and between 8.5 and 10.0 is a high-quality RNA (blue).

#### Quantitative RT-qPCR

The first-strand cDNA was synthesized using the SuperScript IV Reverse Transcriptase (ThermoFisher Scientific, NY, USA) following the manufacturer’s instructions, using 90ng of extracted RNA per sample. Notably, this low quantity of RNA (almost 10 times lower than standard RNA quantity for Real-Time PCR) is due to the limitation of the yield characteristic of LCM collection. Conditions for each cycle of amplification were as follows: heating the RNA-primer mix 5 min at 65° C; after combining the RT reaction mix incubate at 10 min at 55° C, 10 min at 80° C. The final cDNA products were amplified using SyBR Green/ROX PCR Mix (QIAGEN, CA, USA) in 10 μL of a reaction mixture, following the manufacturer’s protocol, using previously designed primers for mouse glyceraldehyde-3-phosphate-dehydrogenase (GAPDH) housekeeping gene (forward, 5’-CCGCCTGGAGAAACCTGCCAAGTAT-3’, reverse, 5’-TTGCTCAGTGTCCTTGCTGGGGT-3’), BDNF (forward: 5’-CTTTGGGGCAGACGAGAAAG-3’, reverse: 5’-TCTCACCTGGTGGAACTCAG-3’), and primers AB from control RNA from cDNA synthesis kit. Real-time PCR was performed using a two-step cycling program, with an initial single cycle of 95° C for 10 min, followed by 40 cycles of 95° C for 15 seconds, then 60° C for 1 min, in an ABI ViiA 7 Real-Time PCR System (Applied Biosystems, NY, USA) with Sequence Detector System software v1.2. A first derivative dissociation curve was built (95°C for 1 min, 65°C for 2 min, then ramped from 65° C to 95° C at a rate of 2° C/min). Non-template control (NTC) was included in the plate to eliminate the possibility of PCR contamination. RNA electrophoretic traces indicates the absence of RNA degradation and high quality, as shown in the two peaks (markers, 18S, and 28S ribosomal RNA; fig 5A). The amplification acquired by RT-qPCR shows a confident expression of the GAPDH gene that is inside the assurable range (cycle threshold>12), according to MIQE (Bustin, 2009; Schroeder, 2006; fig 5B). As expected, we found an opposite association of Ct values from all of 70 samples and RIN. Furthermore, to quantify these results obtained by Agilent Bioanalyser and Ct values, we analyzed the normalized expression of the Brain Derived Neurotrophic Factor (BDNF) gene by ΔCt showing PCR amplification among all samples (Fig 5C).

**Figure 5.**
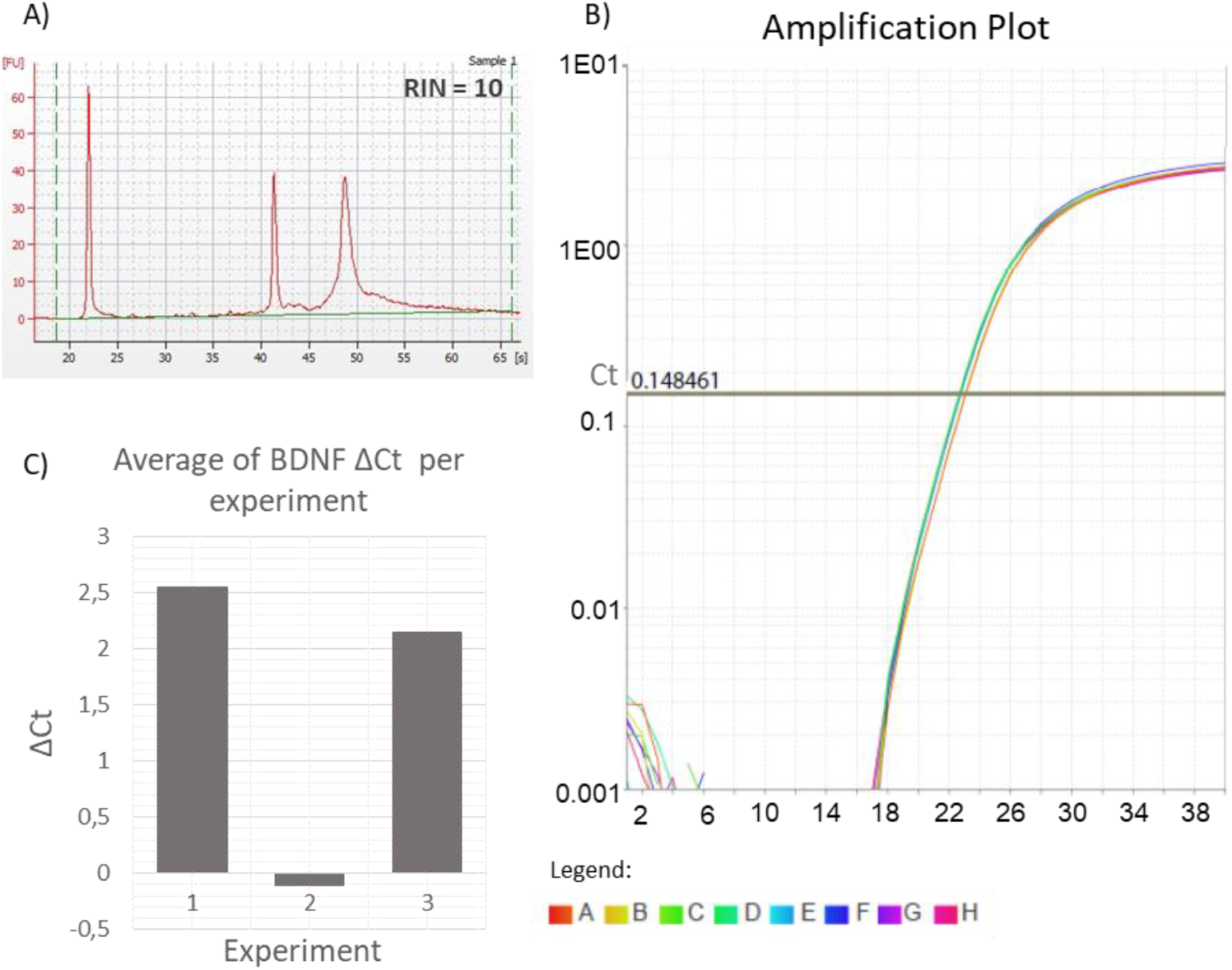
RNA quality verification and validation. **(A)** Electropherogram and electrophoresis assay from representative of anterior cingulate cortex RNA (RIN = 10). **(B)** Amplication plot of housekeeping gene using LCM sample. qPCR detecting valid expression (Ct between 18 and 19) of the housekeeping gene GAPDH in RNA isolated from LCM collected dentate gyrus from nine mice. cDNA from non-template control was used as a negative control. Delta Rn corresponds to normalized reporter SYBR by passive reference dye minus the baseline fluorescence signal. The threshold line was automatically defined as 0,148461. **(C)** Mean BDNF ΔCt values per experiment.

Accurate genetic data from LCM samples are implicated in preserving the good quality of genetic molecules. In the case of a brain microdissected sample, RNA quality depends critically on the time between freezing tissue, the temperature of manipulation, proper slides preparation and staining, and choosing an adequate method of RNA extraction and isolation. Our protocol shows a high percentage of high-quality RNA. We, therefore, conclude that our optimized protocol can be useful for experiments using microdissected mouse brain tissue, resulting in a high degree of purity of RNA.

### Time Considerations

In order to prevent degradation, time managing is crucial. LCM is a powerful tool, however, a disadvantage of the method is that it is time-consuming (S et al., 2000), especially in our approach, in which we prepare slides and collect the samples on the same day. It takes one day long of work to collect brain tissue from one single animal. Experiments with several animals and a control group may take several days to be concluded. However, it is worth ensuring high-quality RNA. If initial procedures are done beforehand, the basic protocol can be completed in a single day, for three different regions from one brain. The cleaning process and stabilizing the brain at −20°C takes at least 30 min. Each slide should take ∼1 hr to cryosectioning, staining, and dissecting. We recommend preparing around 8 slides per day containing 6 slices. This quantity can vary depending on the expected amount. 2 minutes to dissect the brain, 30 minutes to stabilize the brain at −20 °C, 2-3 minutes to the slide dry in the cryostat, 2 minutes in the 70% ethanol RNase free, 30 seconds in violet cresyl, a short dip in 70% ethanol RNase free, a short dip in 100% ethanol RNase free, 1-2 minutes to dry at room temperature, ∼ 30 minutes to cut and catapult. Temperature is important too. To ensure high RNA quality, we strongly recommend keeping all the solutions cold. If all this time is strictly followed, there will be great chances to get optimal results.

## Conflicts of Interest

The authors report no conflict of interest in this article.

## Notes

### Competing Interest Statement

The authors have declared no competing interest.

## Literature Cited

Bustin, S. A., Benes, V., Garson, J. A., Hellemans, J., Huggett, J., Kubista, M., Mueller, R., Nolan, T., Pfaffl, M. W., Shipley, G. L., et al. 2009. The MIQE guidelines: minimum information for publication of quantitative real-time PCR experiments. Clinical Chemistry 55:611–622.

Chabrat, A., Doucet-Beaupré, H., and Lévesque, M. 2015. RNA Isolation from Cell Specific Subpopulations Using Laser-capture Microdissection Combined with Rapid Immunolabeling. Journal of Visualized Experiments: JoVE.

Chung, S. H., and Shen, W. 2015. Laser capture microdissection: from its principle to applications in research on neurodegeneration. Neural Regeneration Research 10:897–898.

Cummings, M., Mappa, G., and Orsi, N. M. 2018. Laser Capture Microdissection and Isolation of High-Quality RNA from Frozen Endometrial Tissue. Methods in Molecular Biology (Clifton, N.J.) 1723:155–166.

Datta, S., Malhotra, L., Dickerson, R., Chaffee, S., Sen, C. K., and Roy, S. 2015. Laser capture microdissection: Big data from small samples. Histology and histopathology 30:1255–1269.

Erickson, H. S., Albert, P. S., Gillespie, J. W., Rodriguez-Canales, J., Marston Linehan, W., Pinto, P. A., Chuaqui, R. F., and Emmert-Buck, M. R. 2009. Quantitative RT-PCR gene expression analysis of laser microdissected tissue samples. Nature Protocols 4:902–922.

Fend, F., and Raffeld, M. 2000. Laser capture microdissection in pathology. Journal of Clinical Pathology 53:666–672.

Gallagher, S. R. 2011. Quantitation of DNA and RNA with absorption and fluorescence spectroscopy. Current Protocols in Neuroscience Appendix 1:Appendix 1K.

Garrido-Gil, P., Fernandez-Rodríguez, P., Rodríguez-Pallares, J., and Labandeira-Garcia, J. L. 2017. Laser capture microdissection protocol for gene expression analysis in the brain. Histochemistry and Cell Biology 148:299–311.

Gilbrich-Wille, C. 2013. History of Laser Microdissection. Available at: https://www.leica-microsystems.com/science-lab/history-of-laser-microdissection/ [Accessed February 25, 2020].

Mahalingam, M. 2018. Laser Capture Microdissection: Insights into Methods and Applications. Methods in Molecular Biology (Clifton, N.J.) 1723:1–17.

Mauney, S. A., Woo, T.-U. W., and Sonntag, K. C. 2018. Cell Type-Specific Laser Capture Microdissection for Gene Expression Profiling in the Human Brain. Methods in Molecular Biology (Clifton, N.J.) 1723:203–221.

Meda, S., Freund, N., Norman, K. J., Thompson, B. S., Sonntag, K.-C., and Andersen, S. L. 2019. The use of laser capture microdissection to identify specific pathways and mechanisms involved in impulsive choice in rats. Heliyon 5:e02254.

Morrison, J. A., Bailey, C. M., and Kulesa, P. M. 2012. Gene profiling in the avian embryo using laser capture microdissection and RT-qPCR. Cold Spring Harbor Protocols 2012.

Mueller, O., Hahnenberger, K., Dittmann, M., Yee, H., Dubrow, R., Nagle, R., and Ilsley, D. 2000. A microfluidic system for high-speed reproducible DNA sizing and quantitation. Electrophoresis 21:128–134.

Nichterwitz, S., Benitez, J. A., Hoogstraaten, R., Deng, Q., and Hedlund, E. 2018. LCM-Seq: A Method for Spatial Transcriptomic Profiling Using Laser Capture Microdissection Coupled with PolyA-Based RNA Sequencing. Methods in Molecular Biology (Clifton, N.J.) 1649:95–110.

PALM MicroBeam Available at: https://www.zeiss.com/microscopy/int/products/laser-microdissection/microbeam.html [Accessed February 25, 2020].

S, C., Ja, M., Hl, M., and Gi, M. 2000. Laser capture microscopy. Molecular Pathology:MP 53:64–68.

Schaeck, M., De Spiegelaere, W., De Craene, J., Van den Broeck, W., De Spiegeleer, B., Burvenich, C., Haesebrouck, F., and Decostere, A. 2016. Laser capture microdissection of intestinal tissue from sea bass larvae using an optimized RNA integrity assay and validated reference genes. Scientific Reports 6:21092.

Schroeder, A., Mueller, O., Stocker, S., Salowsky, R., Leiber, M., Gassmann, M., Lightfoot, S., Menzel, W., Granzow, M., and Ragg, T. 2006. The RIN: an RNA integrity number for assigning integrity values to RNA measurements. BMC Molecular Biology 7:3.

Sonntag, K.-C., and Woo, T.-U. W. 2018. Laser microdissection and gene expression profiling in the human postmortem brain. Handbook of Clinical Neurology 150:263–272.

Thermo Fisher Scientific, T. F. S. I. 2010. Nucleic Acid. Thermo Scientific NanoDrop Spectrophotometers. Available at: https://assets.thermofisher.com/TFS-Assets/CAD/manuals/ts-nanodrop-nucleicacid-olv-r2.pdf.

